# PEG10 viral aspartic protease domain is essential for the maintenance of fetal capillary structure in the mouse placenta

**DOI:** 10.1101/2021.03.02.433660

**Authors:** Hirosuke Shiura, Ryuichi Ono, Saori Tachibana, Takashi Kohda, Tomoko Kaneko-Ishino, Fumitoshi Ishino

## Abstract

The therian-specific gene paternally expressed 10 (*Peg10*) plays an essential role in placenta formation: *Peg10* knockout (KO) mice exhibit early embryonic lethality due to severe placental defects. The PEG10 protein exhibits homology to long terminal repeat (LTR) retrotransposon GAG and POL proteins, therefore mice harboring a mutation in its highly conserved viral aspartic protease motif in the POL-like region were generated because it is essential for LTR retrotransposons/retroviruses. Intriguingly, frequent perinatal lethality, not early embryonic lethality, was observed with fetal and placental growth retardation starting mid-gestation. In the mutant placentas, severe defects were observed in the fetal vasculature, where PEG10 is expressed in the three trophoblast cell layers that surround fetal capillary endothelial cells. Thus, *Peg10* has essential roles not only in early placenta formation, but also in placental vasculature maintenance from mid- to late-gestation. This implies that along the feto-maternal placenta interface an interaction occurs between two retrovirus-derived genes, *Peg10* and retrotransposon Gag like 1 (*Rtl1,* also called *Peg11*), that is essential for the maintenance of fetal capillary endothelial cells.

**Summary statement:** Disruption of the highly conserved viral aspartic protease domain in PEG10 causes placental abnormality leading to perinatal lethality in mice.

## Introduction

*PEG10* is a therian-specific gene encoding GAG- and POL-like proteins that exhibit at most 20~30 % homology with long terminal repeat (LTR) sushi-ichi retrotransposon (Ono et al., 2001). Certain characteristic traits, such as the presence of a CCHC RNA-binding domain in the GAG-like region, a DSG viral aspartic protease motif in the POL-like region and a −1 ribosomal frameshift that results in a PEG10-open reading frame (ORF)1 and an ORF2 fusion protein (PEG10-ORF1/2), are highly conserved among all of the eutherian and marsupial orthologues of PEG10 (Shigemoto et al., 2001; Clark et al., 2007; Suzuki et al., 2007) (Fig. S1), although *PEG10* has lost the LTR sequences at both ends of its structure. These unique features, together with its emergence in therian mammals, indicate that *PEG10* was domesticated in a common therian ancestor from an extinct retrovirus approximately 166 million years ago (Suzuki et al., 2007; Warren et al., 2008). We previously demonstrated that *Peg10* KO mice exhibit early embryonic lethality before 10.5 days post coitus (dpc) due to severe placental dysplasia, including the loss of trophoblast cells in the labyrinth and spongiotrophoblast layers (Ono et al., 2006). The labyrinth layer is an essential part of the mouse placenta in which nutrient and gas exchange occurs between the fetal and maternal blood, and trophoblast cells are unique to the placenta (Rossant and Cross, 2001). These features imply that the acquisition of *PEG10* was a critical event in the emergence of the viviparous reproduction system in the common therian ancestor(s) (Kaneko-Ishino and Ishino, 2015).

In what manner is *PEG10* involved in placenta development? It has been reported that *PEG10* is involved in the development and progression of cancer (Okabe et al., 2003; Tsou et al., 2003; Li et al., 2006). Unfortunately, the biochemical functions of the PEG10 protein have remained obscure since its discovery in 2001, possibly because of the range of its structural/functional varieties due to the properties of the GAG and POL proteins. GAG and the GAG-POL fusion proteins are digested by the aspartic protease activity in POL to form several distinct parts, each with a specific role/function. These include three structural proteins, matrix, capsid and nucleocapsid proteins from GAG along with four enzymatic proteins, aspartic protease, reverse transcriptase, RNase H and DNA integrase from POL (Kirchner and Sandmeyer, 1993; Dunn et al., 2002; Pettit et al., 2004). Similarly, PEG10 is reported to be digested by the highly conserved aspartic protease in the Pol-like ORF2 (Clark et al., 2007). Therefore, it is conceivable that PEG10 plays multiple roles during the course of development, not only in the placenta, but also in other organs and tissues. Which part and/or motif of the PEG10 is essential for early placenta formation associated with trophoblast differentiation and growth, and what are the functions of other parts and/or motifs of PEG10? It is clear that a systematic approach is necessary to solve this puzzle in a step-by-step manner, so a study was undertaken to generate a series of *Peg10* mutant mice harboring a mutation for each of the highly conserved traits. In this study, we analyzed *Peg10-*protease-motif mutant mice and found that PEG10 is expressed in the trophoblast cell layers at the feto-maternal interface and the mice underwent perinatal lethality due to severe defects in feto-maternal circulation in the placenta. This result clearly shows that *Peg10* plays different roles in placental development in a stage-dependent manner, placenta formation via trophoblast differentiation and growth in the early gestational stage and maintenance of the fetal capillary network from mid- to late-gestation, and that PEG10 protease activity is indispensable for the latter process.

Interestingly, retrotransposon Gag like 1 (*RTL1*, also called *PEG11*), another paternally expressed imprinted gene, exhibits homology to the same sushi-ichi retrotransposon as *PEG10* (Charlier et al., 2001; Seitz et al., 2003; Youngson et al., 2005). It is a eutherian-specific gene and plays an essential role in the maintenance of the fetal capillary network in mid- to late-gestation on the endothelial cell side (Sekita et al., 2008; Kitazawa et al., 2017). Thus, it is likely that there exists some interaction between the two retrovirus-derived genes *Peg10* and *Rtl1* along with their effects at the feto-maternal interface in the mouse placenta for maintaining the fetal vasculature, via their functions in trophoblast and endothelial cells, respectively.

## Results and Discussion

### Disruption of PEG10 DSG protease activity leads to both fetal and placental growth retardation, resulting in frequent perinatal lethality

The DSG protease domain in the Pol-like PEG10-ORF2 is highly conserved among therian mammals (Fig. S1), suggesting the importance of the function(s) of this domain. In an effort to determine what function the PEG10 DSG protease plays in placental development, we generated a mutant strain harboring a point mutation in the DSG domain by replacing aspartic acid (D) with an alanine (A) residue using the CRIPSR-Cas9 system (Fig. 1A). It is reported that the PEG10 DSG protease exerts self-cleavage activity so as to produce several fragments and that the same amino acid substitution (DSG to ASG) disrupts the protease activity of the human PEG10 protein *in vitro* (Clark et al., 2007). As *Peg10* is a paternally expressed gene, mice carrying a paternally transmitted mutant allele (+/ASG) (hereafter called *Peg10*-ASG mice) were used for phenotypic analysis throughout this study. First, we examined whether *Peg10*-ASG mice exhibit the early embryonic lethality like *Peg10* null KO mice (+/−). Unexpectedly, the mutant embryos and placenta exhibited an apparently normal appearance at 12.5 dpc (Fig. S2). On western blotting experiments (Fig. 1B), a PEG10 self-cleavage product (approximately 75 kDa in this figure) was observed only in the wild-type (+/+) placenta, it was undetectable in the *Peg10*-ASG placenta, thus confirming the fact that the mutation from DSG to ASG disrupts its protease activity, as expected. These results demonstrate that the loss of the DSG protease function is not responsible for the early embryonic lethality associated with the differentiation and growth defects of placental trophoblast cells observed in *Peg10* null KO mice.

**Figure 1.**
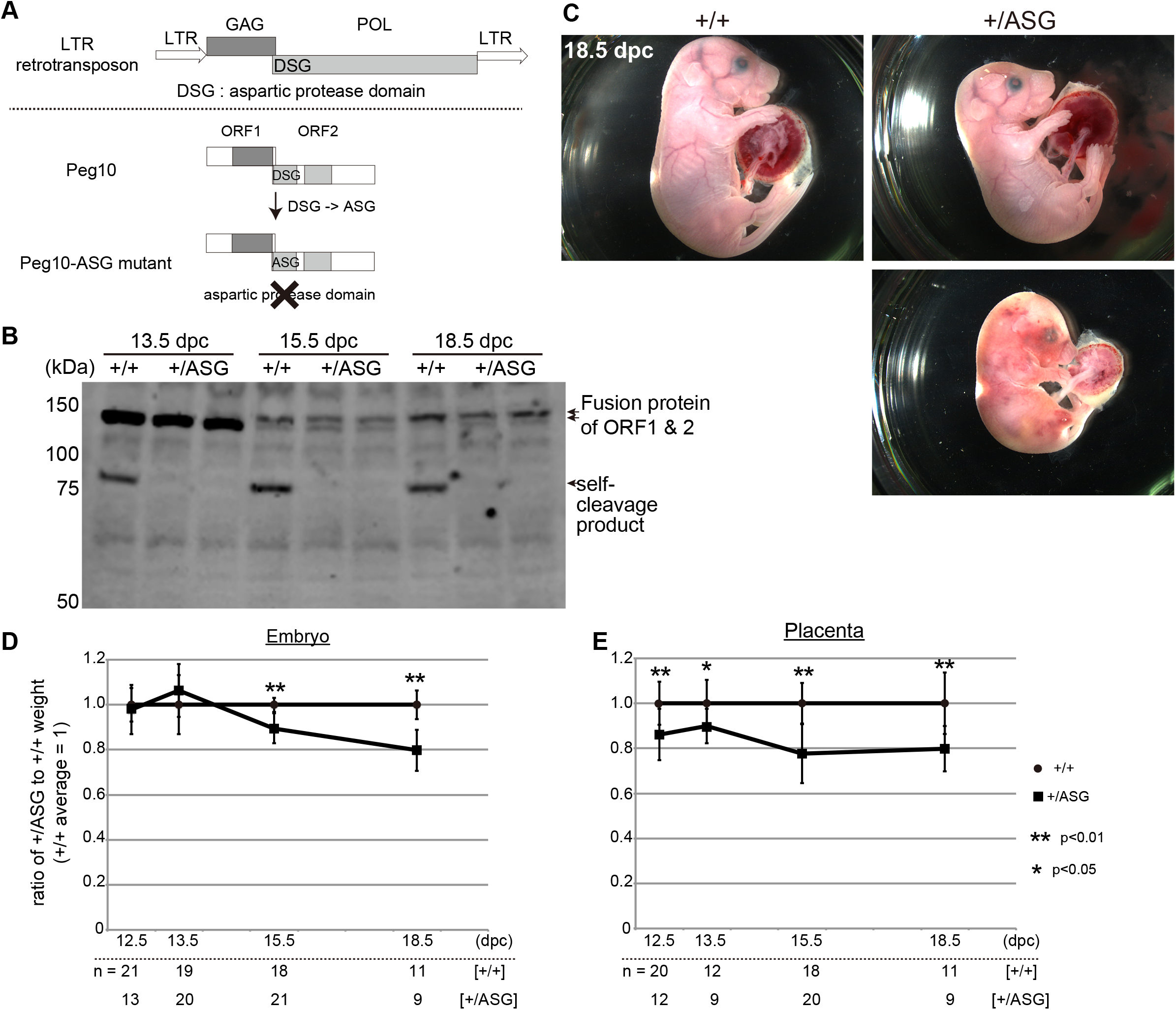
Generation of *Peg10*-ASG mice harboring the PEG10 protease mutation. (A) Schematic representation of the *Peg10*-ASG mutation. The aspartyl protease domain composed of DSG residues was disrupted by substitution of alanine (A) for the aspartic acid (D) residue. (B) PEG10 protein expression in placenta was confirmed by using an anti-PEG10-ORF2 antibody. PEG10 self-cleaved products (approximately 75 kDa) were undetectable in the +/ASG placentas. (C) Wild type (+/+) (top left) and +/ASG (top right) embryos at 18.5 dpc. Some of the +/ASG embryos recovered at this stage were already dead (bottom right). (D) (E) Growth curves of the +/+ and +/ASG embryos (D) and the placental weight (E). Mean weights were calculated for each genotype within a given litter and a value of 1 represents the mean weight of the +/+ mice. The mean values ± SD of each genotype were plotted (* : p < 0.05, ** : p < 0.01).

At the weaning stage, the total rate of mutant pups was extremely low, only 14% (5 mutants/35 pups including both wild-types and mutants) compared with the expected value of 50%(Table 1), indicating that most of the *Peg10*-ASG pups died during the course of perinatal development and growth. Approximately 40 % of the recovered mutant fetuses were found dead at 18.5 dpc (10 dead fetuses/26 total fetuses) (Fig. 1C and Table 1). Moreover, the surviving mutant fetuses were of small size and had small placentas compared to normal mice (Fig. 1C). The mutant embryos were of normal weight before 13.5 dpc, but their growth retardation started around 15.5 dpc and reached ~20% reduction at 18.5 dpc (Fig. 1D), while the placental weight reduction was already evident at 12.5 dpc and persisted to term (Fig. 1E). These results demonstrate that the DSG protease activity is essential for placental growth in mid- to late-gestation and that the inactive ASG mutation apparently caused placental hypoplasia, which consequently led to fetal growth retardation and perinatal lethality.

**Table 1.**
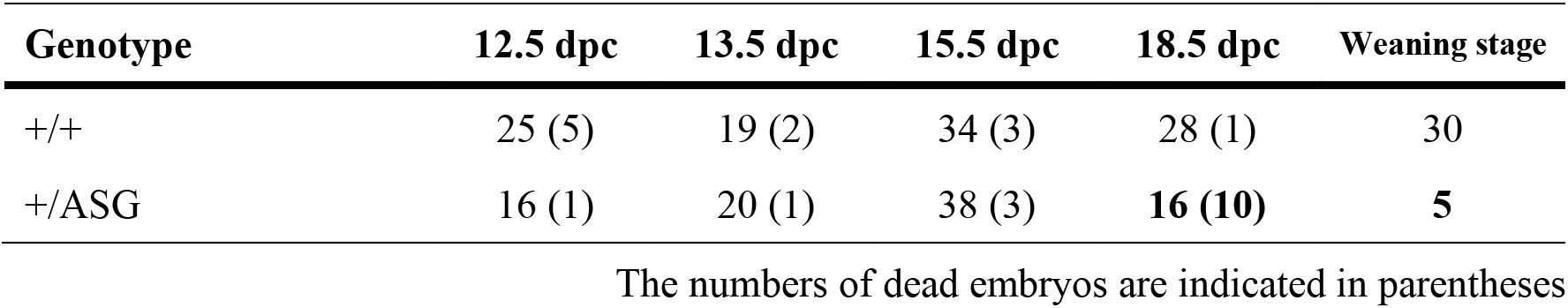
The number of embryos or pups from +/+ dams crossed with +/ASG or ASG/+ sires.

### *Peg10*-ASG mice exhibited severe defects in the vascular network in the labyrinth layer

Evident morphological abnormalities were observed in the labyrinth layers of the *Peg10*-ASG placenta. The fine, mesh-like structure of the fetal vascular network was severely damaged. The fetal vasculature is comprised of fetal capillary endothelial cells and three surrounding layers of trophoblast cells, two syncytiotrophoblast (SynT-I and II) layers and one mononuclear trophoblast layer. On sections of the 18.5 dpc placenta, alkaline phosphatase (AP) staining was employed to distinguish the maternal blood sinus (Adamson et al., 2002) (Fig. 2A), while the immunohistochemistry used for the endothelial marker CD31 (Fig. 2B) revealed severe endothelial cell deformation and collapse of most of the fetal capillaries in the labyrinth layers, resulting in a loss of the fine, mesh-like structure of the fetal vasculature. We performed simultaneous immunofluorescence-staining using CD31 and a pan-trophoblast marker cytokeratin (CK) (Fig. 2C) to confirm a physical relationship between the fetal endothelial cells and trophoblast cells. In the normal placenta, the endothelial cells and trophoblast layers closely aligned along the fetal capillaries. In contrast, this alignment became irregular and their respective locations separated from each other in the *Peg10*-ASG placenta. Under magnified views, the fetal endothelial cell nuclei were observed to have accumulated in narrow spaces and as a result the fetal blood spaces had become clogged (Fig. 2D). Taken together, these results suggest that abnormal organization of the fetal vasculature causes severe impairment of feto-maternal exchange in the labyrinth layer, leading to the perinatal lethality of the *Peg10*-ASG fetuses and pups.

**Figure 2.**
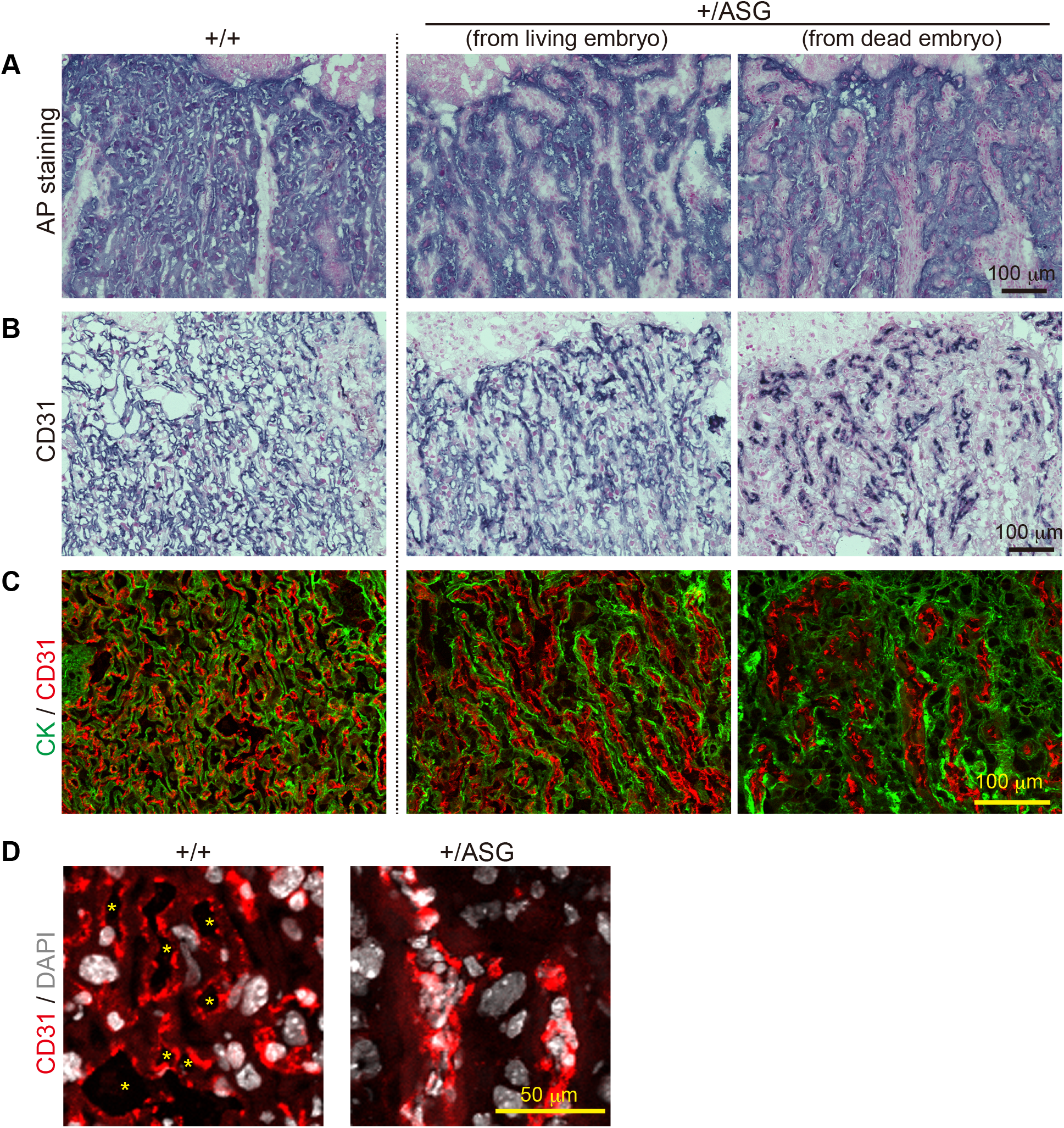
Abnormal vascular organization in the +/ASG placentas. (A) (B) Trophoblast cells (A) and fetal endothelial cells (B) in the labyrinth layers at 18.5 dpc were detected by alkaline phosphatase (AP) staining and immunohistochemical staining with an anti-CD31 antibody, respectively. Nuclei were stained with Nuclear Fast Red. (C) Immunofluorescence analysis of the 18.5 dpc labyrinth layers with an anti-cytokeratin (CK) (trophoblast cells marker, green) and CD31 (red) antibodies. (D) Magnified images of immunofluorescence with an anti-CD31 antibody (red) and nuclear staining with DAPI (white). The asterisks indicate fetal blood spaces.

### Loss of DSG protease results in fibrinoid necrosis-like lesions in the placental labyrinth layer

Next, placental morphology was assayed by Hematoxylin and Eosin staining (Fig. 3A). We found that the *Peg10*-ASG labyrinth layer in dead embryos was deeply stained in shades of pink-red and contained considerable endothelial cell nuclear debris, while the remaining area stained in bright pink contained enlarged trophoblast cell nuclei. This staining pattern is characteristically seen in one of the necrotic forms of cell death, fibrinoid necrosis (Damjanov, 2009). Additionally, immunostaining for the pan-leukocyte marker CD45 revealed that the number of leukocytes was markedly increased in *Peg10*-ASG placentas obtained from both living (data not shown) and deceased embryos (Fig. 3B). These results suggest that severe inflammation was induced in the fetal vasculature of the *Peg10*-ASG placenta, which then led to the development of fibrinoid necrosis-like lesions in the labyrinth layers, ultimately resulting in severe fetal damage and/or death.

**Figure 3.**
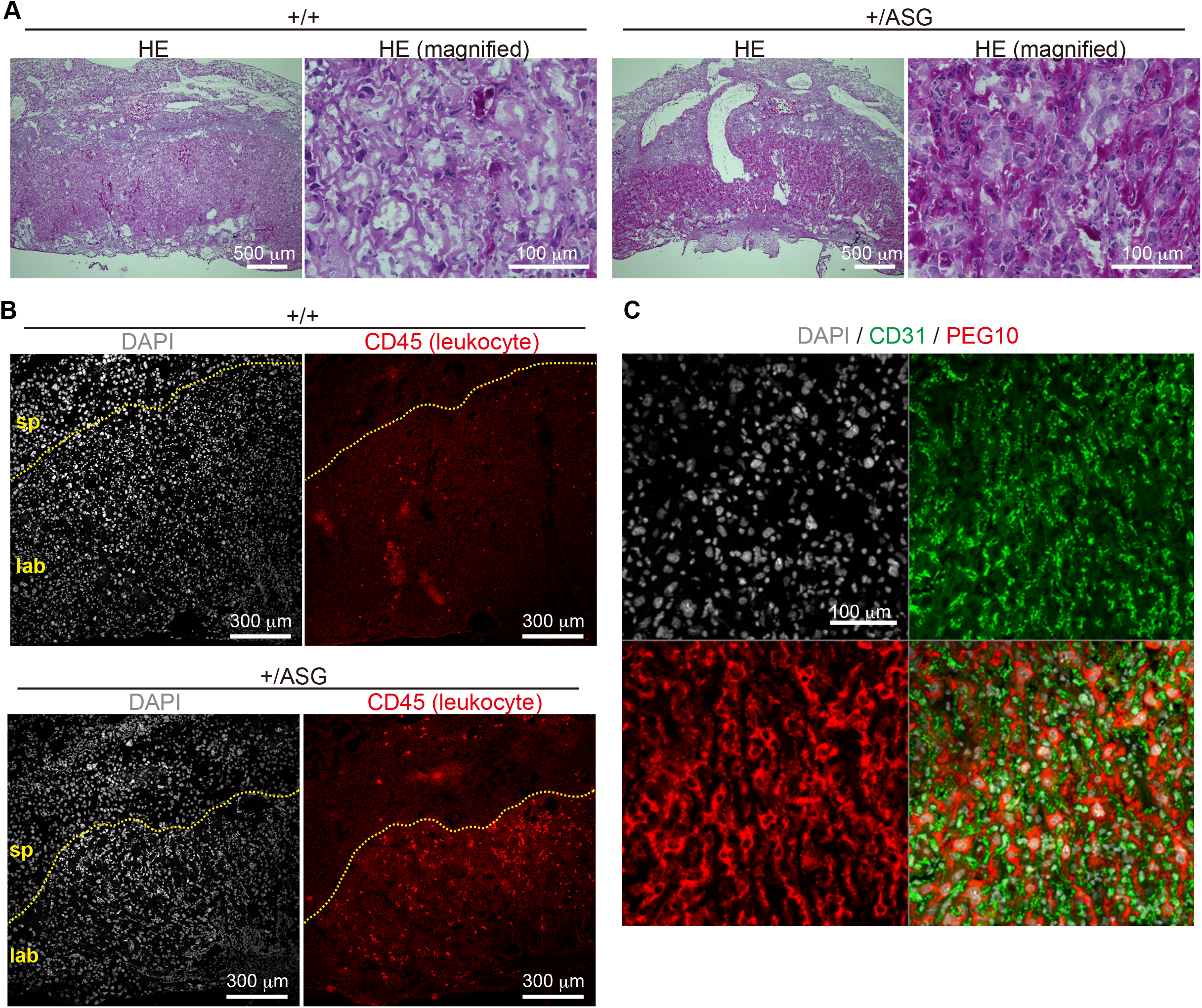
The +/ASG placental labyrinth layers exhibit fibrinoid necrosis-like lesions. (A) HE-stained histological sections of the 18.5 dpc placenta. Low and high magnification images of each +/+ and +/ASG placenta are represented side-by-side. (B) Immunofluorescence analysis of the 18.5 dpc placentas with an anti-CD45 antibody (pan-leucocyte marker, red). Nuclei were stained with DAPI (white). sp and lab indicate spongiotrophoblast and labyrinth layers, respectively. (C) Immunofluorescence analysis of the 18.5 dpc wild-type placenta with an anti-PEG10 (red) and CD31 (green) antibodies. Nuclei were stained with DAPI (white).

### Interaction between two retrovirus-derived genes at the feto-maternal interface in the eutherian placenta

In this study, it was demonstrated that the disruption of the protease activity of the DSG domain in PEG10 results in perinatal lethality associated with fetal and placental growth retardation, probably because of the severe damage inflicted on the fetal vascular network. It is likely that such extensive fetal capillary damage would result in a disturbance of the feto-maternal circulation in the placental labyrinth. This indicates that the viral-type aspartic protease domain in PEG10 is essential during the mid to late stage of placental development, implying the protease activity that is intrinsic to retrovirus/LTR retrotransposon was successfully domesticated (or exapted) to the placental functions in eutherians.

Importantly, as shown in Fig. 3C and Fig. S3, the PEG10 protein exhibited high expression in the three trophoblast cell layers but no localization in the CD31-positive fetal endothelial cells that were also damaged in the *Peg10*-ASG placentas. It is thus strongly suggested that the trophoblast cells in the fetal capillary have some defense mechanism to protect the fetal endothelial cells against non-specific hazardous events and that direct or indirect effects of PEG10 DSG protease activity is largely involved in this process. We have demonstrated in previous studies that *Rtl1* (or *Peg11*), which is derived from the presumable same retrovirus as *Peg10*, is essential for the maintenance of the fetal capillaries in the placenta (Sekita et al., 2008; Kitazawa et al., 2017). It should be noted that, in contrast to *Peg10*, *Rtl1* is only expressed in endothelial cells (Sekita et al., 2008). This demonstrates that the two retrovirus-derived genes *Peg10* and *Rtl1* function on either side of the feto-maternal interface, i.e. in trophoblasts and endothelial cells in the fetal capillaries, respectively, and presumably act in a cooperative manner so as to maintain a normal fetal vasculature for the exchange of nutrients/waste and O_2_/CO_2_ gas between the fetal and maternal blood (Fig. S4). Recent phylogenic analysis has revealed that the mouse and human type of hemochorial placentas, which have a feto-maternal interface in which the trophoblast surface has direct contact with the maternal blood, is ancestral eutherian placenta (Wildman et al., 2006; Roberts et al., 2016). Therefore, it is highly likely that domestication of *PEG10* and *RTL1,* before and after divergence of the eutherians and marsupials, respectively, must have been critical events and exerted a driving force in the evolution of the eutherian viviparous reproduction system via their effect on the eutherian placenta.

*PEG10* and *RTL1* are expressed in many other tissues and organs besides the placenta (Abed et al., 2019; Kitazawa et al., 2021). Recently, we demonstrated that mouse *Rtl1* plays other important roles during the course of development, such as in fetal/neonatal skeletal muscle maturation as well as in functional activity of the central nervus system (Kitazawa et al., 2020; Kitazawa et al., 2021). Therefore, it is possible that *PEG10* has a multiplicity of functions across a range of organs and tissues. Future studies are needed to show how *PEG10* contributed to the evolution/acquisition of various other therian-specific traits, and it may turn out both *PEG10* and *RTL1* were domesticated in the same process and function cooperatively in the current eutherian developmental systems, as in the case of the maintenance of the fetal vascular network in the placenta.

## Materials and methods

### Mice

All animals and experimental procedures were approved by the Animal Ethics Committees of Tokyo Medical and Dental University and the University of Yamanashi. Animals were allowed access to a standard chow diet and water *ad libitum* and were housed in a pathogen-free barrier facility with a 12L:12D cycle.

### Plasmid preparation

The plasmids expressing *hCas9* and sgRNA were prepared by ligating oligos into the BbsI site of pX330 (http://www.addgene.org/42230/). The 20 bp sgRNA recognition sequences are shown below.

*Peg10*-ORF2-sgRNA (5′-GTCCGAGCTATGATTGATTC-3′)

The oligo DNAs for co-injection with the plasmid into mouse zygotes are shown below. ACCTGCAAGTGATGCTCCAGATTCATATGCCGGGCAGACCCACCCTGTTTGTCCGAG CTATGATTG**C**TTCTGGTGCATCTGGCAACTTCATTGATCAAGACTTTGTCATACAAAA TGCAATTCCTCTCAGAAT

### Production of *hCas9* mRNA and *Peg10*-ORF1-sgRNA

To produce the *Cas9* mRNA, the T7 promoter was added to the *Cas9* coding region of the pX330 plasmid by PCR amplification, as previously reported (Wang et al., 2013). The T7-*Cas9* PCR product was gel purified and used as the template for *in vitro* transcription (IVT) using the mMESSAGE mMACHINE T7 ULTRA kit (Life Technologies). The T7 promoter was added to the *Peg10*-ORF2-sgRNA region of the pX330 plasmid by PCR purification using the primers listed below.

*Peg10*-ORF2-IVT-F (TTAATACGACTCACTATAGGTCCGAGCTATGATTGATTC), IVT-R (AAAAGCACCGACTCGGTGCC)

The T7-sgRNA PCR product was gel-purified and employed as the template for IVT using the MEGAshortscript T7 kit (Life Technologies). Both the *Cas9* mRNA and *Peg10*-ORF2-sgRNA were DNase-treated to eliminate template DNA, purified using the MEGAclear kit (Life Technologies), and eluted into RNase-free water.

### Generation of Peg10-ASG mice

B6D2F1 female mice were superovulated and IVF carried out using B6D2F1 mouse sperm. The synthesized hCas9 mRNA (50 ng/µl) and *Peg10*-ORF2-sgRNA (25 ng/μl) with oligo DNA (10 ng/μl) were injected into the cytoplasm of fertilized eggs at the indicated concentration. The eggs were cultivated in KSOM overnight, then transferred into the oviducts of pseudopregnant ICR females.

### Preparation of frozen sections

The recovered mouse placentas were directly embedded in OCT compound (Sakura Finetek) or first fixed in 4 % paraformaldehyde, incubated sequentially in 10%/15% sucrose in PBS for 2 hr at 4℃, and 25 % sucrose in PBS overnight at 4℃, and ultimately embedded in OCT compound (Sakura Finetek). The OCT blocks were sectioned at a thickness of 7 μm with a cryostat (MICROTOME) and mounted on Superfrost Micro Slides (Matsunami Glass).

### Histological analysis

The frozen sections were fixed in 4 % PFA for 10 min at room temperature and washed three times with PBS for 5 min. The sections were then subjected to hematoxylin and eosin (HE) staining, immunodetection (immunofluorescence and immunohistochemistry) or endogenous alkaline phosphatase activity detection.

For HE staining, the sections were stained with HE, and mounted with malinol mounting medium (Muto Pure Chemicals).

For immunodetection, after incubation in blocking buffer (1% BSA in PBS with 0.1% TritonX-100) for 60 min, the sections were reacted with the primary antibody (for CD31, 1:100 rat monoclonal anti-CD31 (BD Biosciences, 550274); for Cytokeratin, 1:200 rabbit polyclonal anti-Cytokeratin (DAKO, Z0622); for CD45, 1:100 rat monoclonal anti-CD45 (BD Biosciences, 561037); for PEG10, 1:200 rat monoclonal anti-PEG10-ORF1 (produced in-house) in blocking buffer at 4°C overnight. After washing with PBS three times for 5 min each, the sections were incubated with a secondary antibody conjugated with a biotin or fluorescence label (for immunofluorescence) or biotin for immunohistochemistry (for CD31, PEG10 and CD45, 1:500 goat anti-rat IgG conjugated with Cy3 (Thermo Fisher Scientific, A10522); for Cytokeratin, 1:500 goat anti-rabbit IgG conjugated with Alexa 488 (Thermo Fisher Scientific, A11008)) in blocking buffer at 4°C overnight. For immunofluorescence, after washing with PBS three times for 5 min each, the slides were mounted with VCTASHIELD-Hardset (Vector Laboratories) containing 1 μg/ml DAPI. The sections were imaged under confocal fluorescence microscopy (LSM510; Carl Zeiss). For immunohistochemistry, after washing with PBS three times for 5 min each, the slides were incubated with ABC-AP reagent (VECTASTAIN ABC-AP Kit (Vector Laboratories)) with 200 nM levamisole for 30 min at room temperature. The sides were washed in PBS, followed by incubation with 100 mM Tris-HCl (pH 9.5) for 10 min at room temperature. Chromogenic detection was performed by NBT-BCIP staining (BCIP/NBT AP substrate (Vector Laboratories)) according to the manufacturer’s instructions. After rinsing with 100 mM Tris-HCl (pH 9.5), the sections were counterstained with Nuclear Fast Red (Vector Laboratories).

For endogenous alkaline phosphatase activity detection, after being incubated with 100 mM Tris-HCl (pH 9.5) for 10min at room temperature, chromogenic detection was performed as described above.

### Western blot analysis

Placenta tissue lysates were resolved by SDS–PAGE and transferred to a PVDF membrane (BioRad) before immunoblotting. The membrane was then probed with the rabbit polyclonal anti-Peg10-ORF2 antibody (produced in house), followed by incubation with horseradish peroxidase-conjugated donkey anti-rabbit IgG (GE Healthcare). The blots were developed using the ECL system according to the manufacturer’s instructions (Thermo Fisher Scientific).

## Acknowledgements

We thank T. Usami (MRI, TMDU) for generation of *Peg10-*ASG mice. Pacific Edit reviewed the manuscript prior to submission.

## Competing interests

The authors declare that no competing interests exist.

## Funding

This work was supported by the funding program for Grants-in-Aid for Scientific Research (A) (16H02478 and 19H00978) to F. I., Grants-in-Aid for Scientific Research (C) (26430183) to R. O. from Japan Society for the Promotion of Science (JSPS), Nanken-Kyoten (Grant No.2020-40), Tokyo Medical and Dental University (TMDU) to H.S. and F.I. and The Mochida Memorial Foundation for Medical and Pharmaceutical Research to H.S.

## Notes

### Competing Interest Statement

The authors have declared no competing interest.

